# Absolute Quantification of Aging-Associated Glycans in IgG for Biological Age Prediction: Insights from Glycomics and Transcriptomics

**DOI:** 10.1101/2025.01.05.631349

**Authors:** Huijuan Zhao, Jiteng Fan, Jing Han, Wenjun Qin, Jichen Sha, Weilong Zhang, Yong Gu, Xiaonan Ma, Jianxin Gu, Shifang Ren

## Abstract

Immunoglobulin G (IgG) N-glycans have been identified as associated with aging; however, previous studies predominantly quantified changes based on the relative percentages of each glycan within the total glycan pool, neglecting the absolute concentration changes of individual glycans. Additionally, relative quantification can limit practical applications, as these often require absolute concentration measurements for consistent and interpretable biomarker values. In this study, we introduce a novel strategy for discovering aging-associated IgG glycans and establishing a prediction model based on their absolute concentration alteration. We employed a glycome quantification technology to identify the alteration in IgG glycan amount in natural aging and anti-aging (caloric restriction) models, discovering aging-related glycans. The glycomics analysis revealed key features: downregulation of bisected glycan GP3 (F(6)A2B) and upregulation of digalactosylated glycan GP8 (F(6)A2G2). These glycan changes showed significant fold changes from an early stage. Using external standards of these two glycans, we subsequently measured the absolute concentrations of them, allowing us to establish a predictive model, abGlycoAge, for biological aging. The abGlycoAge index suggested a younger state under caloric restriction, with an average age reduction of 3.9-14 weeks. Additionally, we performed RNA sequencing on splenic B cells of mice, suggesting Derl3, Smarcb1, Ankrd55, Tbkbp1 and Slc38a10 could contribute to the alteration of GP3 and GP8 in aging. This analysis enhances our understanding of glycan alterations, accounts for individual variability, and aids in designing effective anti-aging strategies. These findings highlight the crucial roles of the GP3 and GP8 as potential biomarkers for aging and health.

## INTRODUCTION

Aging induces complex physiological changes across multiple systems, making it essential to identify reliable biomarkers that can quantify these changes, explore the underlying mechanisms, and support the development of targeted interventions^1^. Among these changes, immunosenescence is a well-recognized feature, which involves the gradual decline and imbalance of immune function, increasing the risk of infections, chronic inflammation, and cancer^2,3^. Recent research has focused on molecular biomarkers that reflect immunosenescence, including alterations in glycosylation patterns, especially on immunoglobulin G (IgG), as these changes have been linked to heightened inflammatory responses and immune dysfunction ^4^. Studies have shown that IgG accumulates during aging in mice and humans^5,6^. What’s more, quantification of IgG N-glycans has shown promise in assessing biological age, with studies suggesting potential causal links between specific glycan structures and aging ^7-11^. These findings demonstrate the potential of glycosylation as a biomarker for biological age and as a tool to explore age-related immune changes and guide therapeutic interventions.

However, while IgG N-glycans analysis has contributed significantly to understanding aging processes, current quantification techniques face limitations in fully defining aging-related glycan changes. Previous studies have primarily detected IgG N-glycans using high-performance liquid chromatography (HPLC) coupled with fluorescence detection, which quantifies each glycan as a proportion of the total glycan pool ^9,10,12,13^. The relative quantification approach has been effective in identifying major changes across samples and has led to key discoveries, including aging-associated glycans that have shown promise as biomarker. However, using ratio values for assessment can introduce variability, complicating consistent interpretation across different samples and conditions ^14^. In practical applications, absolute concentration measurements are often preferred, as they enable clear, standardized reference values suitable for diverse applications^15-18^. Furthermore, relative quantification may not adequately capture lower-abundance glycans that, while minor, may have biologically significant in aging^14^. Therefore, absolute quantification^19,20^ offers a complementary approach that enhances the applicability of glycan biomarkers by providing precise, direct measurements that reveal both major and subtle glycan changes.

Mass spectrometry (MS) is widely used for glycomic analysis due to its sensitivity, speed and rich structural information through tandem MS approaches. However, the absolute quantification of glycans, particularly with Matrix-Assisted Laser Desorption/Ionization Time-of-Flight Mass Spectrometry (MALDI-TOF-MS), remains challenging due to issues such as non-linear signal response and the difficulty of finding suitable internal standards for diverse glycan structures^21^. Previous studies reported effective methods by emplorying three N-Glycans and Malto-Series Oligosaccharide Standards or chemo-enzymatic synthesis of ^13^C labeled N-glycan libraries to improve the accuracy of Glycan Quantification by MALDI-MS Analysis ^19^. Our lab previously developed a simple method by one-step reducing to obtain internal standards (Bionic Glycome) with full coverage of the natural N-glycome to be analyzed^22^. This method provides a unique solution to the challenges of glycan quantification by producing a structurally similar internal standard for each glycan, matched in abundance. This tailored internal standard approach minimizes interference from more abundant glycans, ensuring accurate detection even for low-abundance glycans that are often difficult to quantify with conventional methods. Moreover, this method enables assess the amount alteration of each glycan, rather than relative proportion in total glycan pool, allowing it to reveal absolute concentration changes across large sample sets. This capability is particularly valuable for identifying specific aging-associated glycans within complex glycan profiles. Once these specific glycans are identified, a standard curve can be established with external standard glycan using the same MALDI-MS method, leveraging its speed and efficiency. By addressing the limitations of traditional MALDI-MS quantification, the approach in this study enables the development of biomarker models with precise, absolute measurements, which are essential for practical and reproducible biomarker applications.

In this study, we applied ‘Bionic Glycome’ method to analyzed IgG N-glycans of C57BL/6 mice across different age groups, aiming to discover aging-associated glycans. Caloric restriction (CR) was used as an anti-aging intervention to further confirm the association of these glycans with aging. Subsequently, we developed an absolute quantification method for aging-related glycans by using external glycan standards, enabling us to measure absolute concentration changes in these glycans across age and intervention groups. The predictive value of the model instructed with the absolute concentration in practical applications was also evaluated. To gain further insights, we integrated our glycan analysis with transcriptome analysis to examine gene expression patterns associated during aging. By comparing data from natural aging and anti-aging interventions, we identified several genes correlated with aging-related glycan changes, providing insights into future mechanistic studies. This integrative approach not only enhances the utility of glycan biomarkers, but also support the development of predictive biomarkers and potential intervention strategies.

## RESULTS

### 1. Age-related Changes in IgG N-glycosylation Patterns in C57BL/6 Mice

IgG N-glycans from the Aging cohort mouse sample (n=89, 5 per sex at each time point) were analyzed by MALDI-MS based on Bionic Glycome method. The Study design is illustrated in **Figure 1**. A total of 20 glycans were observed, all of which had been previously identified by HPLC-FLR-ESI-TOF MS^23^ or MALDI-TOF-MS^24^ with coefficient of variation (CV) less than 20% in the reproducibility test (**Supplementary Table 1**). Additionally, four glycans (GP10+GP11, GP13, GP15, and GP16) that contained both α2,3- and α2,6-linked sialic acids were combined in the subsequent quantitative calculations to ensure comparability with previous HPLC results^23^ (**Supplementary Table 1**).

**Figure 1.**
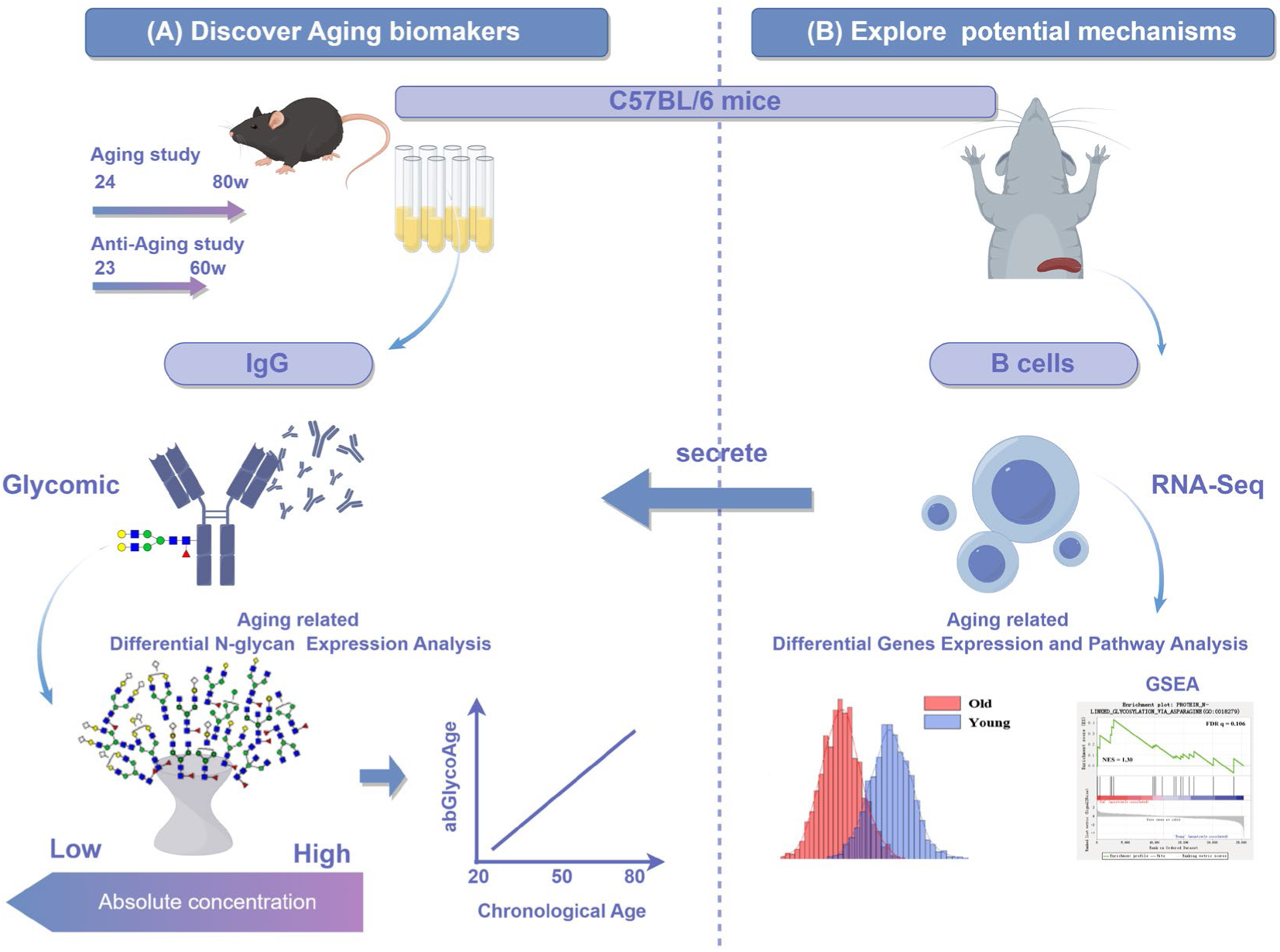
Study design for predicting biological age from aging-associated glycans in IgG by integrating glycomics and transcriptomics. We performed a comprehensive IgG N-glycome profiling in C57BL/6 mice at 7 time points across 80 weeks by MALDI-TOF-MS. Alterations in glycan amount were detected using an Internal Full-Glycome Standard. By referencing Exogenous Standards, we introduced a novel method for detecting aging biomarkers based on their absolute concentration. Based on the absolute concentrations of aging-related IgG glycans, we established abGlycoAge index to evaluate the biological age and applied in Anti-Aging Assessment. The potential molecular basis of these IgG aging-related glycans was explored by splenic B cell transcriptome.

The results revealed a general trend of IgG N-glycosylation increasing early in life, followed by a decline during aging (**Figure 2 A**). In male mice, IgG N-glycosylation levels significantly declined by 68 weeks, whereas in female mice, this decline became prominent at 80 weeks. Correlation analysis indicated sex-specific differences in age-related glycan changes: in males, six glycans (GP1, GP4+GP5, GP7, GP8, GP9 and GP14) were positively correlated with age, while GP3 was negatively correlated. In females, three glycans (GP4+GP5, GP8, and GP14) were positively correlated with age, while four (GP2, GP3, GP6, and GP10+GP11) were negatively correlated **(Figures 2B and 2C)**. When compared to 24-week-old mice, both male and female mice showed significant fold changes in GP3 and GP8 at multiple time points **(Figure 2D)**. Based on this, we identified the key aging-associated glycans in C57BL/6 mice as the downregulation of GP3 (F(6)A2B) and the upregulation of GP8 (F(6)A2G2) **(Figure 2E).**

**Figure 2.**
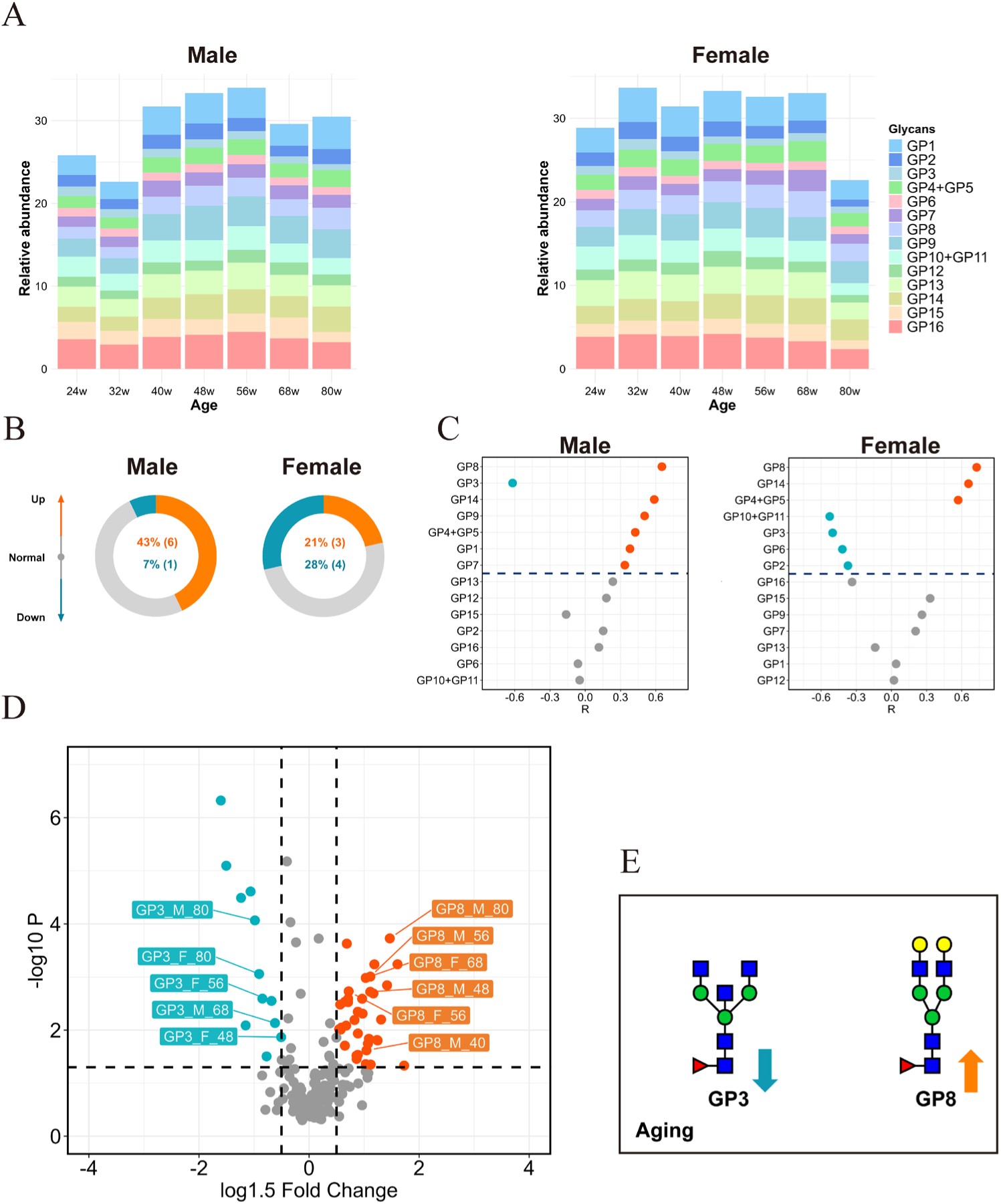
Age-related Changes in IgG N-glycosylation Patterns in C57BL/6 Mice at seven time points (24w, 32w, 40w, 48w, 56w, 68w, 80w). (A) Stacked Bar Chart illustrating the Age-Related Changes in IgG Total N-Glycomes (each color representing a distinct glycan). Schematic comparison of glycan alteration in aging (B and C). Circles show the percentage of glycans that are positively correlated with age (Orange), negatively correlated with age (Green), and no correlation (Gray). (C) Dot plot show the correlation of each glycan with age. (D) Vocano plot show the fold changes (log1.5FC) of glycans expression compared to 24-week-old mice. The differences among various age groups were assessed using the One-Way ANOVA, followed by Benjamini-Hochberg (BH) test procedure to adjust the p-values for multiple comparisons. Adjusted P values <0.05 was considered to be statistically significant. (E) Schematic diagram highlight the crucial roles of the GP3 and GP8 as potential biomarkers for aging and health.

### 2. Caloric Restriction Reverses Key Age-related IgG Glycan Changes in C57BL/6 Mice

We then performed quantitative analysis of individual IgG N-glycans in Caloric restriction (CR) and ad libitum (AL). Notably, the changes in GP3 diminished after long-term CR, likely due to its low relative abundance. Quantitative analysis revealed that GP3 exhibited the most pronounced changes during CR (**Figure 3B&C**). GP3 levels increased significantly in the early stages of CR, and although they gradually decreased with aging, they remained higher than in age-matched AL mice after long-term CR. Additionally, glycans containing LacNAc structures, such as GP4+GP5 and GP8, displayed opposite trends after CR. Compared to the AL group, GP4+GP5 and GP8 were significantly downregulated in the early stages of CR (15-35 weeks) as aging progressed, though this difference diminished slightly over time, yet remained statistically significant **(Figure 3B&C)**. In summary, caloric restriction effectively reverses the age-related downregulation of GP3 (F(6)A2B) and the upregulation of GP8 (F(6)A2G2) in C57BL/6 mice.

**Figure 3.**
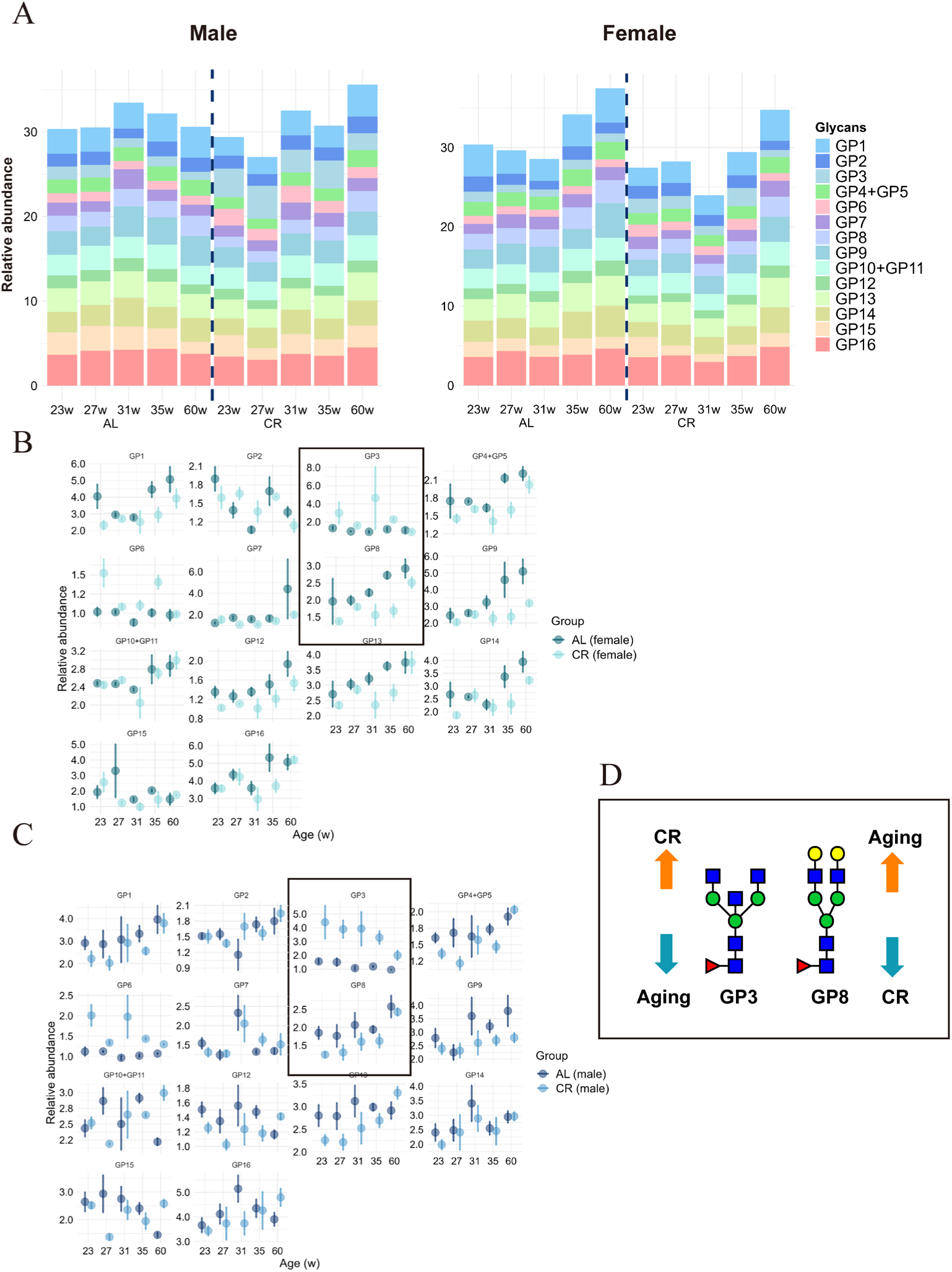
Caloric Restriction Reverses Key Age-related IgG Glycan Changes in C57BL/6 Mice. (A) Stacked Bar Chart illustrate the Age-Related Changes in IgG Total N-Glycomes both in AL and CR group (each color representing a distinct glycan). Schematic comparison of glycan alteration in female (B) and male (C) in AL and CR groups (Mean±SE). (D) Schematic diagram highlight the crucial roles of the GP3 and GP8 as potential biomarkers for aging.

We also detect changes using High-Performance Liquid Chromatography (HPLC) to make a comparison with previous studies. We observed significant changes in the composition of 14 IgG glycans during the early stages of CR (15-35 weeks). Specifically, four N-glycans—GP2, GP3, GP6, and GP10+GP11—showed steady increases across all age groups, while GP4+GP5, GP8, and GP13 steadily decreased. Further analysis of IgG glycan expression in the serum of 60-week-old CR and ad libitum (AL) mice revealed that the changes in GP2, GP3, and GP13 had disappeared, with the CR group showing similar levels to the AL group. However, glycans containing LacNAc structures, such as GP4+GP5, GP8, and GP10+GP11, maintained their early-stage trends, continuing to either decrease or increase (**Supplementary** Figure 1).

### 3. Development of a Glycan-Based Biological Age Prediction Model

In both male and female mice, GP3 and GP8 showed a significant correlation with age, while the correlation between these two glycans was relatively weak (|r| < 0.5). Based on this, we aimed to develop a glycan-based biological age prediction model using GP3 and GP8. First, we performed absolute quantification of GP3 and GP8 ratio using standard glycans. Both glycans exhibited linear standard curves across the sample concentration range, with correlation coefficients (r²) exceeding 0.999. The absolute concentration ranges were 1-25 nM for GP3 and 100-500 nM for GP8 (**Figure 4A a-b**). Using these linear standard curves, we calculated the glycan content per gram of IgG for each mouse (**Figure 4B**) and fitted model for predicting biological age based on the absolute glycan content in mice of different ages (**Figure 4C**).

**Figure 4.**
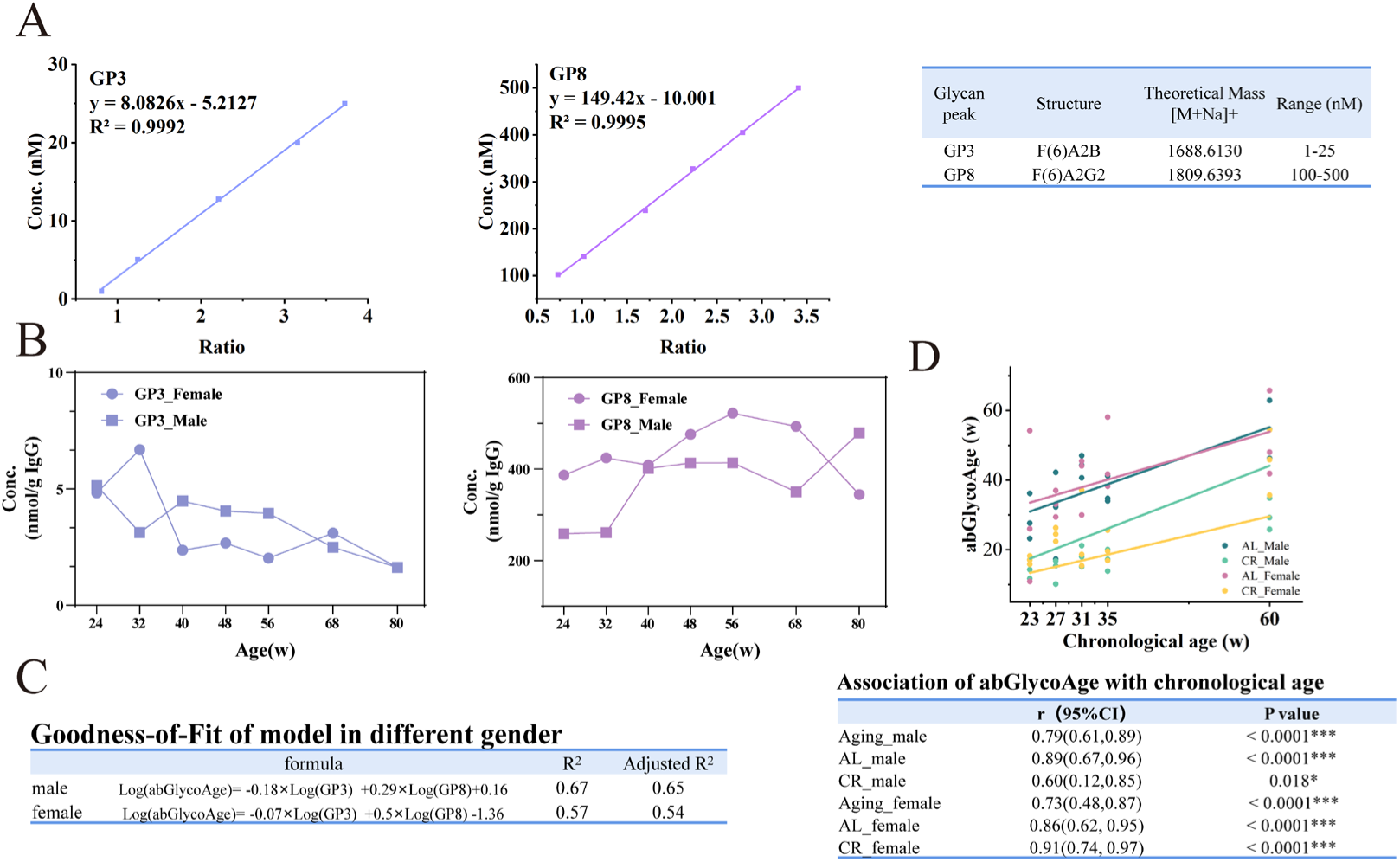
Development the Glycan-Based Biological Age Prediction Model and Assessment Biomarker Sensitivity in Anti-Aging study. (A) Absolute quantification of GP3 and GP8 was performed using standard glycans. The linear standard curves for both glycans exhibited correlation coefficients (r²) greater than 0.999 across the sample concentration ranges of 1-25 nM for GP3 and 100-500 nM for GP8. (B) Glycan content per gram of IgG was calculated for mice in each age group based on the standard curves. (C) A model was fitted to predict biological age (abGlycoAge) using the absolute glycan content in mice of different ages. The model explained 67% of the actual age variance in male mice and 57% in female mice, with a correlation of 0.79 in males and 0.73 in females. (D) The model was applied to CR cohorts and predicted a reduction in biological age (abGlycoAge) by an average of 14.0 weeks in males and 3.9 weeks in females, a reduction sustained throughout the lifespan.

Both GP3 and GP8 were strongly correlated with age in male and female mice, and their correlation with each other was weak (|r|<0.5). Based on these glycan profiles, we developed a biological age prediction model, abGlycoAge, using the absolute quantification of GP3 and GP8. The model explained 67% of the actual age variance in male mice and 57% in female mice, with a correlation of 0.79 in males and 0.73 in females. When applied to predict biological age in CR cohorts, the model estimated that CR reduced biological age by an average of 14.0 weeks in males and 3.9 weeks in females, a reduction maintained throughout the lifespan (**Figure 4D**).

### 4. Mechanistic Exploration: Transcriptomic Analysis of Splenic B Cells

A total of 7,440 differentially expressed genes (DEGs) were observed in the comparisons among three groups. Compared to young mice, 3,114 DEGs were observed in aged mice, with 1,782 upregulated and 1,332 downregulated. Gene set enrichment analysis (GSEA) revealed significant enrichment of the "protein N-linked glycosylation via asparagine" pathway (GO:0018279) in aged mice, suggesting that protein N-glycosylation is upregulated during aging, and this effect is mitigated by CR (**Figure 5A.a-c**).

**Figure 5.**
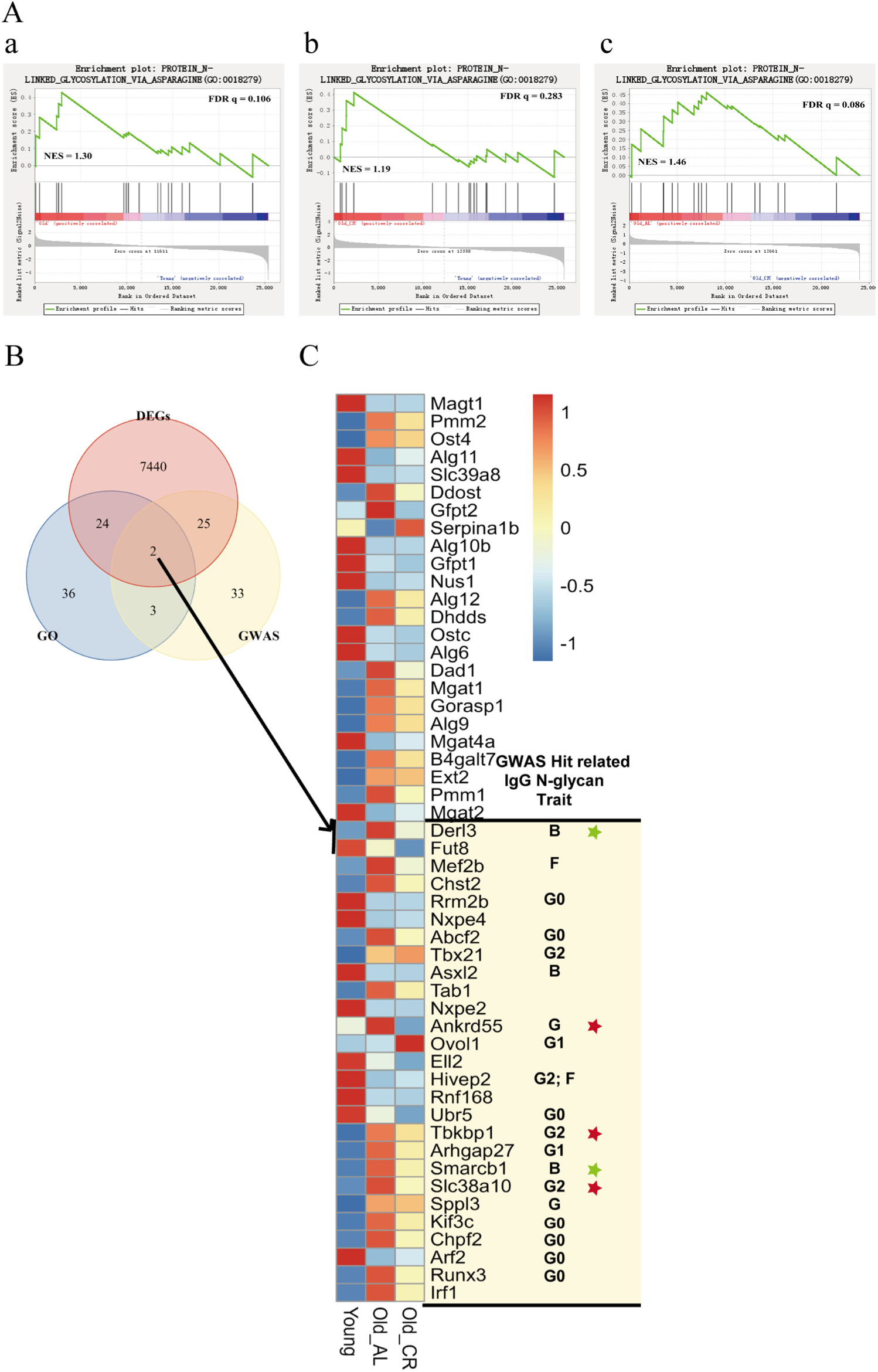
IgG N-Glycan-related genes and pathway perturbation as potential mechanisms underlying variations in glycosylation phenotypes during aging. A. GSEA plots illustrating the activation of protein N-linked glycosylation via asparagine in aged mice across various comparisons: (a) Old vs. Young, (b) Old_CR vs. Young, and (c) Old_AL vs. Old_CR. B. Venn diagram illustrating the IgG N-glycan-related differentially expressed genes (DEGs) identified through by integration of transcriptome analysis, genome-wide association studies (GWAS), and Gene Ontology (GO) terms. DEGs: Differentially expressed genes identified via transcriptome analysis, encompassing the union of DEGs across all possible combinations of the three groups. GWAS: Genes identified in previous genome-wide association studies. GO: Genes associated with protein N-linked glycosylation (GO:0006487). C. The expression patterns of IgG N-glycan-related DEGs for three group (20w for Young, 60w for Old_AL and Old_CR) was shown in the left heatmap.

IgG N-glycan-related DEGs in the aging process were screened to elucidate their potential mechanisms in the expression level changes of the aging biomarkers GP3 (F(6)A2B) and GP8 (F(6)A2G2). Candidate genes were pinpointed by synthesizing data from RNA-seq analyses, the gene ontology term for protein N-linked glycosylation (GO: 0006487) **(Supplementary Table 2-1)**, and prior GWAS**(Supplementary Table 2-2)**. Although these genes are known to be associated with N-glycosylation and IgG glycans, their expression level changes during aging have not been fully studied. As depicted in **Figure 5B**, among the 51 candidate genes, two genes were common across all three datasets, 24 were found in both the GO and DEGs sets, and 25 were shared between the GWAS and DEGs sets. The two common genes were Derl3 and Fut8. Most of the N-glycans were core fucosylated in mice IgG. Therefore, the change of Fut8 in aging appears to influence level of total N-glycans. The *SMARCB1-DERL3* locus were previously implicated in the level of fucosylated structures with bisecting GlcNAc N-glycans of Human IgG^25^. Our research has extended these findings by demonstrating that the expression of these two genes changes with aging and can be reversed in anti-aging interventions, thereby providing additional evidence for the mechanisms underlying the alterations observed in the GP3 (F(6)A2B). Fut8 is an essential α-1, 6-fucosyltrasferase for promoting core fucosylation. Runx3, have been reported to regulate IgG glycosylation in vitro, were also observed in our study as upregulated in aged mice and were restored in anti-aging mice. Another group of genes were Ankrd55, Tbkbp1 and Slc38a10, previously observed influence the percentage of agalactosylated and digalactosylated structures in total neutral IgG glycans. This may be the basis for the alterations observed in the GP8 (F(6)A2G2). (**Figure 5C)**.

## Discussion

Glycans are valuable as biomarkers due to their stability and ability to integrate genetic, epigenetic, and environmental factors into quantifiable chemical structures, making them ideal for personalized medicine ^26^. Aberrant N-glycosylation modulates IgG effector functions flexibly and dynamically, with aging driving a shift in the IgG N-glycome from an anti-inflammatory to a pro-inflammatory composition^27^. These changes in glycan composition are often reflected in specific glycan levels. Previous studies have demonstrated relative quantification is an effective approach for assessing IgG glycosylation changes. However, examining glycan changes from an absolute quantification perspective is more suitable for the practical application of glycans as biomarkers. In clinical applications, widely used biomarkers such as CA-125 and PSA proteins in tumor screening ^16^, metabolites in newborn screening ^18^, and cyclosporin A (CsA) levels in blood for managing immunosuppression after organ transplantation ^15^ all depend on absolute quantification. This study offers valuable insights into specific glycan shifts associated with aging by focusing on their absolute concentrations. The capacity to measure biological aging through glycans holds significant potential for practical applications, including the evaluation of aging processes and the effectiveness of anti-aging interventions.

To our knowledge, this is the first longitudinal study to provide detailed information on the absolute concentration changes of IgG glycan levels in mice. We employed a simple, rapid and reliable method based on MALDI-TOF-MS to assess the alteration of IgG glycan amount, which significantly improved both experimental efficiency and data accuracy. In addition to natural aging, we included CR as anti-aging intervention, enabling a comprehensive analysis of IgG N-glycosylation patterns and offering new perspectives on anti-aging mechanism. With over 100 samples and multiple time points, our study provides a robust and comprehensive dataset. The large sample size and detailed temporal profiling enhance the reliability of our results, offering valuable insights into glycan changes over time.

In this study, we identified two main N-glycans in C57BL/6 mice serum IgG during 24-80 weeks. The downregulation of GP3 (F(6)A2B) and the upregulation of GP8 (F(6)A2G2) emerge as significant characteristics of aging in these mice. To our knowledge, galactosylation levels in human plasma IgG have shown the strongest association with age^7-9^. Despite species-specific differences leading to divergent trends, in our study, we observed age-related changes in GP8 (F(6)A2G2), a glycan with two galactose, in C57BL/6 mice serum IgG, both in relative^23^ and absolute quantification. These findings highlight GP8 as an important hallmark of aging.

In previous studies, bisecting N-acetylglucosamine (GlcNAc) glycans were detected in both mouse and human IgG^9,10,12,13,24^. However, findings regarding the association between bisecting GlcNAc glycans and aging in mice have been inconsistent, likely due to the limitations of relative quantification methods. These studies primarily employed high-performance liquid chromatography (HPLC) with fluorescence detection, which quantifies glycans based on their relative proportions in the total glycan pool. This relative quantification approach is particularly limited when characterizing low-abundance glycans, such as GP3 (F(6)A2B), as their measurement can be significantly influenced by the presence of high abundance glycans. In contrast, utilizing absolute quantification assessment allows for a more comprehensive understanding of bisecting GlcNAc changes during mouse aging, providing clearer insights into the age-related significance in this glycan. Notably, both of the GP3 and GP8 levels were modulated by calorie restriction. This suggests that calorie restriction may delay the aging process by stabilizing GP3 expression and suppressing GP8. However, further mechanistic studies are needed to determine whether these glycosylation changes in IgG are a cause or a consequence of reduced inflammation following caloric restriction.

The transcriptomic analysis of splenic B Cells was integrated to enhance our understanding the relationship between the observed aberrant specific glycans level and aging. To our knowledge, we have, for the first time, unveiled the RNA-seq landscape of splenic B cells within the context of aging, which includes both the natural aging process and the state following anti-aging interventional measures. Previously GWAS^25,28-31^ identified genetic variants linked to specific glycan traits; however, their impact on gene expression is rarely investigated. Besides, the composition ratios of specific glycan obtained by traditional methods are technically difficult in directly correlating with the expression profiles of genes. Our method demonstrated the changes in glycan amount which can theoretically correlated better with gene expression alteration, favoring gaining insight into the underlying mechanism of glycan alteration.

We have focused on genes related to N-glycan synthesis, as well as the 63 genes associated with IgG glycosylation reported in previous GWAS^25,28-31^. We indeed observed the activation of key pathways regulating the synthesis of IgG N-glycans, through "protein N-linked glycosylation via asparagine" pathway. This finding is consistent with the observed increase in total N-glycan levels in aged mice. Additionally, we have identified changes in the expression levels of 27 out of the 63 genes associated with IgG glycosylation as reported by GWAS, with 3 of these genes (Hivep2, Sppl3, Runx3) also previously validated through in vitro experiments for their correlation with IgG glycan expression levels, which indicates the reliability of our findings. Furthermore, we provided evidence that the expression levels of the other 24 genes may also regulate IgG glycosylation in vivo. We further validated the association of these genes with aging by investigating whether the alteration of gene expression during naturally aging can be reversed in anti-aging model.

Among them, five genes were previously found to be associated with the bisected glycans and galactosylated glycans ^25,31^. The bisecting and galactosylation are also the main traits of GP3 and GP8, respectively. The finding suggested that Derl3 and Smarcb1 might be main possible contributor of the alteration of GP3. Similarly, Ankrd55, Tbkbp1 and Slc38a10 might be main possible contributor of the alteration of GP8. Our research provides their expression characteristics in vivo and, through aging and anti-aging studies, suggests that these genes may constitute the main molecular basis for the IgG glycan GP3 and GP8 during the aging process.

In summary, this study highlights the downregulation of GP3 (F(6)A2B) glycans and the upregulation of GP8 (F(6)A2G2) glycans during the aging process of C57BL/6 mice, elucidates the role of calorie restriction in modulating these glycan changes as part of it anti-aging effects. We also demonstrated the potential of bisected glycans and digalactosylated glycans as biomarkers of aging and health. By establishing a glycan age prediction model, abGlycoAge, this study quantifies the biological relevance of glycosylation in the aging process. Furthermore, the application of absolute quantification techniques has facilitated the precise detection the alterations in low-abundance glycans and improved the feasibility of using discovered specifically changed glycans as practical biomarkers. In addition, our finding indicate a potential novel anti-aging strategy: precisely modulating GP3 and GP8 on IgG through glycoengineering. Together, these findings not only deepen our understanding of glycan changes during aging but also support the development of new anti-aging intervention strategies and predictive biomarkers.

## MATERIALS AND METHODS

### Chemicals

Sodium dodecyl sulfate (SDS), 1-hydroxybenzotriazole monohydrate (HOBt), sodium borodeuteride (NaBD4), trifluoroacetic acid (TFA), sodium hydroxide (NaOH), 2-aminobenzamide (2-AB), super-2,5-dihydroxybenzoic acid (super-DHB) and Sepharose CL-4B, ammonium bicarbonate (ABC) were purchased from Sigma-Aldrich (St. Louis, MO, USA). 1-ethyl-3(3-(dimethylamino)propyl)-carbodiimide (EDC) hydrochloride was purchased from Fluorochem (Hadfield, U.K.). HPLC SupraGradient acetonitrile (ACN), ethanol (EtOH) and formic acid (FA) were provided by Merck (Darmstadt, Germany). Protein G BestaroseTM 4FF beads was purchased from Bestchrom (Shanghai, China). NP-40 and peptide-N-glycosidase F (PNGase F) were from Adamas-life (Shanghai, China). The FiltrEX™ 96-well Clear Filter Plates with 0.2 μ m PVDF Membrane was purchased from Corning (NY, USA). Protein concentrations were determined using the Pierce™ BCA Protein Assay Kit (Thermo Fisher Scientific).

### Anlysis of IgG N-glycans

#### Isolation of IgG from murine serum

IgG was isolated by a high-throughput manner using affinity chromatography as described previously ^23^. In brief, murine IgG was captured from 30 μL serum by Protein G BestaroseTM 4FF beads. Each serum sample was diluted 3-fold with 1 × PBS and incubated with beads for 30 min. The beads were washed with PBS and nano-pure water. IgG was eluted with 100 μL 100 mM FA and neutralized with 1 M ABC. Studies showed that IgG accumulates during aging in mice and humans ^5,6^.To exclude the influence of protein concentration, each sample was adjusted to 25 μg and vacuum dried at room temperature.

#### Preparation of Released IgG N-glycans

Dried murine IgG samples were dissolved in 5 μL water and 10 μL 2% SDS and denatured for 10 min at 60 °C. Then 10 μL glycobuffer (4% Nonidet P-40, 5 × PBS, PH 7.5) and 1 μL of 10-fold diluted PNGase F were added to the mixture followed by an incubation at 37 °C for 16 h.

#### Preparation of N-glycans internal standard

The Bionic Glycome of murine serum IgG was prepared as the internal standard ^22^. Briefly, a mouse serum mix was prepared to isolate IgG and then release N-glycans. Once the volumes of ethanol (EtOH) and 1% formic acid (FA) were added, the mixture was incubated at 37°C for 2 hours. Then, the N-glycan were reduced by 2M NaBD_4_ at 60°C for 2 hours.

#### Glycans enrichment and purification

The Sepharose HILIC solid-phase extraction in 96-well plate format was performed as described previously ^32^. Specifically, N-glycan mixtures with acetonitrile was added to the activated Sepharose beads, and N-glycans were adsorbed to the beads. After three washes with 95% acetonitrile containing 1% TFA, three washes with 95% acetonitrile, retained glycans were eluted with water, vacuum dried and stored at -20°C.

#### Sialic acids derivatization

Prior to MALDI-TOF-MS analysis, the samples were re-dissolved in 6.5 μL water and internal standards were re-dissolved in 10.5 μL water. The sialic acids were stabilized by ethyl esterification ^33^. Two microliters of mixture from sample or internal standards were added with 20 μL esterification reagent (250mM EDC hydrochloride and 250mM HOBt monohydrate in ethanol) and incubated in 37°C for 1 hour. Cotton HILIC SPE microtips were used to microscale purification and enrichment glycans. Finally, N-glycans were eluted in 10 μL water.

#### Quantification of Glycans

The N-glycan standards GP3 (F(6)A2B) and GP8 (F(6)A2G2) were obtained from the laboratory of Professor Tiehai Li (Shanghai Institute of Materia Medica, Chinese Academy of Sciences). First, a series of standards solutions with different concentrations were prepared to ensure coverage of the expected sample concentration range. Subsequently, the ratio of the target glycan to the Bionic Glycome internal standard in each standard solution was measured by MALDI-TOF-MS. These ratios were used to construct a linear standard curve, and the relationship between the target glycan concentration and the ratio was determined through linear regression analysis. Using these linear standard curves, we calculated the target N-glycan concentration in eluate for each sample. Furthermore, we estimated the glycan content per gram of IgG for each mouse using this formula:

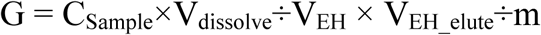

G: the glycan content per gram of IgG, nmol/g. C_Sample_: the N-glycan concentration of sample in eluate, nmol/L. V_EH_: the volume of glycan used to ethyl esterification (2μL). V_EH_elute_: the volume of water to elute the retained glycans (10μL). V_dissolve_: the volume of water to re-dissolved dried glycan (6.5μL). m: the amount of IgG used to analysis, 25μg.

#### MALDI-TOF-MS detection

Bruker UltrafleXtreme laser equipped with Smartbeam-II was used to detect the ethyl esterified IgG N-glycans operated under flexControl (Bruker Daltonics). Each sample was mixed with an equal volume of the internal standard and dot 1 μL on a MALDI target (MTP 384 polished steel BC; Bruker Daltonics), with three duplicate dots per sample. After air-dried, one microliter 5 mg/mL super-DHB in 50% ACN and 1 mM NaOH was added and allow the spots to dry by air. Measure the samples in a reflected positive ion mode. The mass window of m/z was set from 1000 to 3000. For each spectrum, 8,000 laser shots were accumulated at a laser frequency of 1000 Hz, using a complete random walk with 100 shots per raster spot.

#### Data processing

The mass spectra were processed with flexAnalysis v3.4 (Bruker Daltonics). The acquired spectra were internally recalibrated using a set of calibration masses (1485.5337, 1647.5865, 1809.6393, 1982.7081, 2144.7609, 2479.8826) and the Glycan composition were known (**Supplementary Table 1**). Masses were picked in the spectra using a sum algorithm, followed by quadratic calibration. A total of 20 pairs of glycans were identified (S/N>3). The recalibrated spectra were imported into the BioPharma Compass software (Bruker Daltonics) to batch extract peak intensity of each spectrum. The analysis was performed as a target data extraction using a determined list of glycan compositions calculated as [M+ Na]^+^. The relative peak signal intensity (light/heavy peak intensity) of each pair of target glycan was used for quantification, and the interday and intraday repeatability of the glycans were examined. Glycans structures were assigned by GlycoWorkbench (v 2.1). Data analysis and visualization were performed in R (v 4.3.3) using the following package: readxl, dplyr, tidyr, rio, stringr, tidyverse, ggplot2, tidymodels. The differences among various age groups were assessed using the One-Way ANOVA, followed by Benjamini-Hochberg (BH) test procedure to adjust the p-values for multiple comparisons. P value<0.05 was considered to be statistically significant.

### Splenic B cells transcriptome analysis

The mouse CD19+ B cells were isolated from mouse spleen cell suspensions using the B Cell Isolation LS Column, and a MidiMACS™ Separator (Miltenyi, Bergisch Gladbach, Germany). The total RNA was extracted from these isolated B cells. The gene expression profiles were compared among the three young mice (20 w), six aged mice (60 w) with an ad libitum diet, and five aged mice (60 w) with calorie restriction. The library construction and sequencing were performed in the Illumina platform provided by Novogene China (Beijing). DESeq2^34^ package (v1.20.0) was used for identifying differentially expressed genes (DEGs, applying a threshold of | log2(Fold Change)| > 1 and False Discovery Rate (FDR) < 0.05). Gene Set Enrichment Analysis (GSEA) ^35^ was conducted by gsea (v4.3.2) for functional annotation of gene expression profiles. Parameters were set as followed: Permutation type = “permute phenotype”, Metric for ranking genes = “signal2noise”, set_min = 15, set_max = 5000, while all other variables were set as default.

IgG N-glycan-related DEGs were screened by integration of the union of DEGs across all possible combinations of the three groups, the genes included in protein N-linked glycosylation (GO:0006487) retrieved from the Mouse Genome Database (MGD)^36^ and the candidate genes identified by previous GWAS^25,28-31^. The R packages “biomaRt” (2.60.0) was used to convert gene IDs. Plots were produced using Origin 2021 (OriginLab, USA) and R packages “pheatmap” (1.0.12).

## Supporting information

sTable1

sTable2-1

sTable2-2

## Contributions

Huijuan Zhao and Jiteng Fan contributed equally to this work. **Huijuan Zhao**: Experiments, Data Processing and Writing - original draft. **Jiteng Fan**: Sample collection, Experiments, Data Processing, Writing - original draft. **Jing Han**: Sample collection. **Wenjun Qin**: Methodology. **Jichen Sha**: Sample collection. **Weilong Zhang**: Sample collection, Experiments. **Yong Gu**: Sample collection, Experiments. **Xiaonan Ma**: Investigation. **Shifang Ren**: Conceptualization, Methodology, Writing - review & editing. **Jianxin Gu**: Conceptualization, Project administration.

## Acknowledgments

We thank Prof. Tiehai Li from the Shanghai Institute of Materia Medica, Chinese Academy of Sciences, for his support in supplying GP3 and GP8 N-glycan external standards.

This work was supported by thea grant from the National Key R&D Program of China (2022YFC3400803) and the National Natural Science Foundation of China (92478201, 32071276).The graphical study design was drawn by FigDraw (www.figdraw.com).

---

**Supplementary Figure 1.**
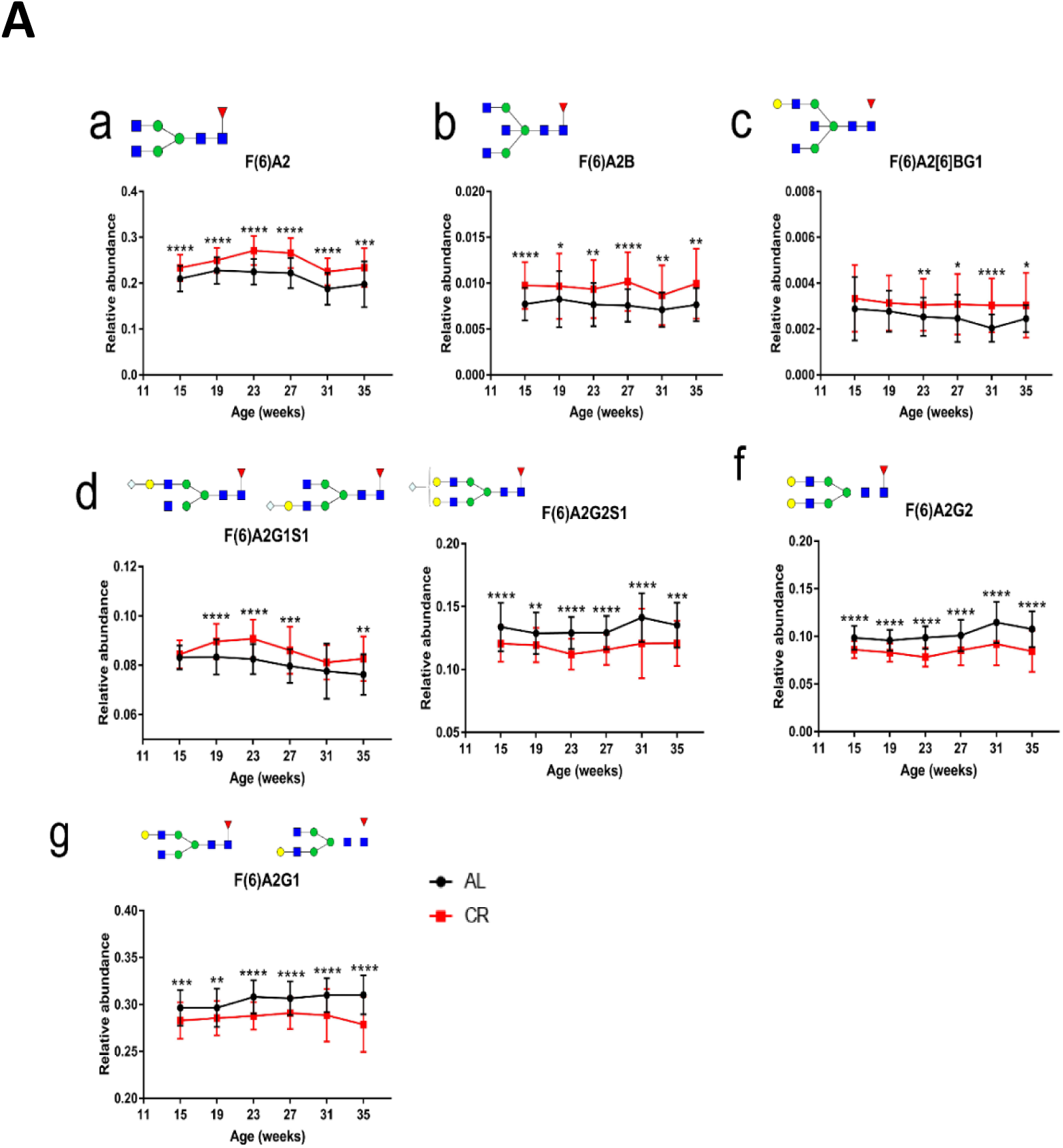
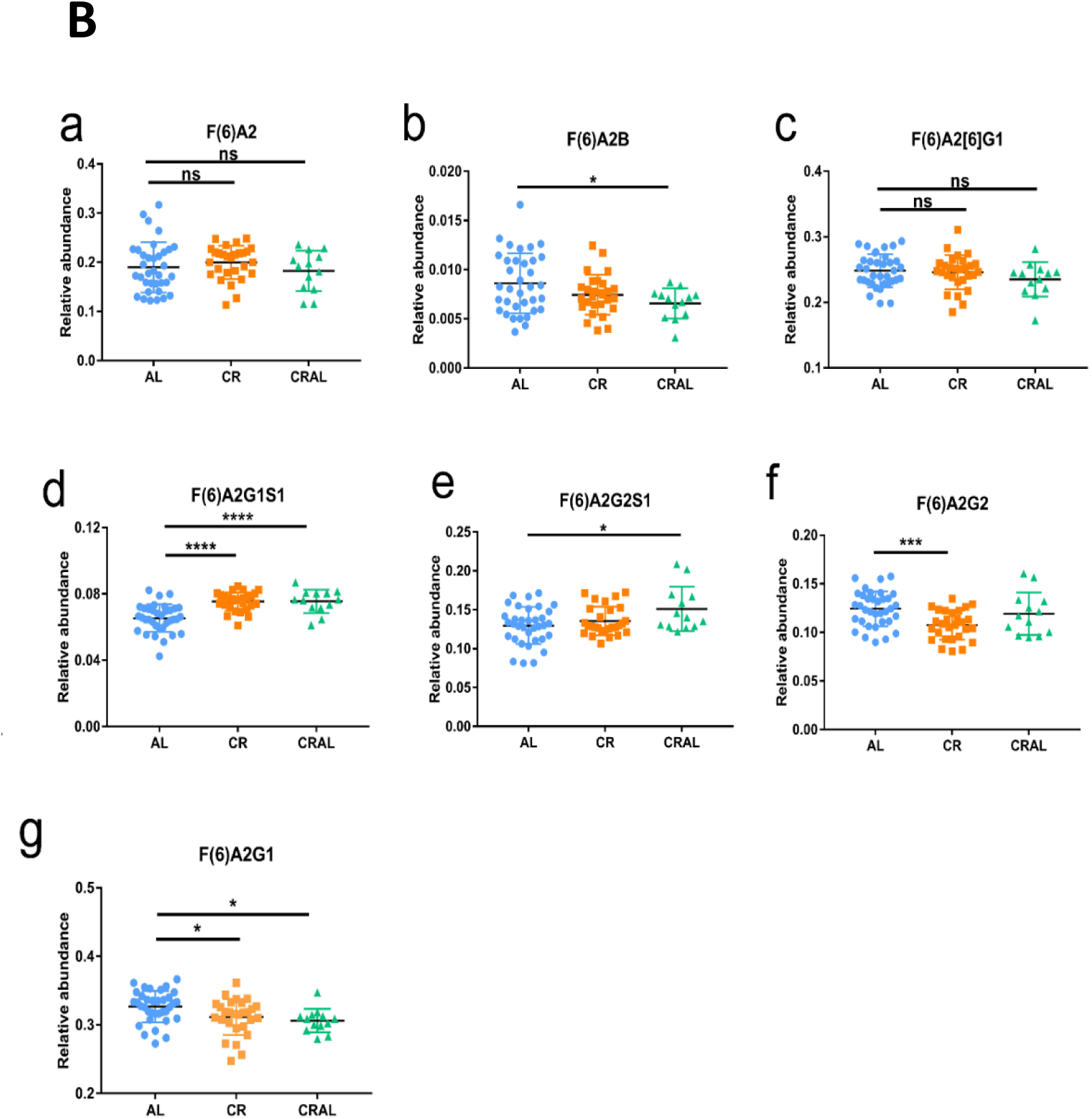
The changes in IgG N-glycans that show significant variations both in the early stages of CR and at the age of 60 weeks in HPLC results. (A) The IgG N-glycans and derived sugar changes that show significant variations in the early stages of CR, and the IgG N-glycans that undergo significant changes after CR. Data are presented as mean ± standard deviation (SD). (B) The changes in some glycans at the age of 60 weeks, and the variations of IgG N-glycans that significantly changed in the early stages of CR among the AL, CR, and CRAL groups at the age of 60 weeks. Structure abbreviations: A1, mono-antennary glycan; A2, diantennary glycan; B, bisected N-acetylglucosamine; F(6), core fucose; Gx, number of β1,4-linked galactose; [6]G, galactose located on the α1,6-mannose antenna;[3]G, galactose located on the α1,3-mannose antenna; G(3), galactose located on the α1,3-galactose antenna; Sx, number of N-acetylneuraminic acids linked to galactose.

