## Supplementary material for "Absolute Quantification of Aging-Associated Glycans in IgG for Biological Age Prediction: Insights from Glycomics and Transcriptomics": sTable1

**sTable 1: The murine IgG N-glycans.**Composition abbreviations: H, hexose; N, HexNAc; F, fucose; Ge, ethyl esterified N-glycolylneuraminic acid (α2,6-linked); Gl, lactonized N-glyco-lylneuraminic acid (α2,3-linked); E, ethyl esterified N-acetylneuraminic acid (α2,6-linked); L, lactonized N-acetylneuraminic acid (α2,3-linked). Structure abbreviations: A1, mono-antennary glycan; A2, diantennary glycan; B, bisected N-acetylglucosamine; F(6), core fucose; Gx, number of β1,4-linked galactose; [6]G, galactose located on the α1,6-mannose antenna;[3]G, galactose located on the α1,3-mannose antenna; G(3), galactose located on the α1,3-galactose antenna; Sx, number of N-acetylneuraminic acids linked to galactose.

| Detected by UPLC-Fluorescence | | Detected by MALDI-TOF-MS | | | | | | | | | | |
|---|---|---|---|---|---|---|---|---|---|---|---|---|
| | | | | Observed in | | Quantified in | Theoretical m/z [M+Na]+ | | CV % Intraday (n=3) | | | Interday (n=9) |
| Glycan peak from UPLC-Fluorescence ( Han et al., 2020) | Structure | Composition | Possible stucture matching the composition | Reference study ( Manfred et al., 2017) | This study | This study | Sample | Internal Standard | day-1 | day-2 | day-3 | |
| GP1 | F(6)A1 | H3N3F1 | 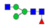 | √ | √ | √ | 1282.4543 | 1285.4762 | 5.10 | 2.88 | 9.01 | 6.23 |
| GP7 | M6 | H6N2 | 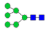 | × | √ | √ | 1419.4755 | 1422.4974 | 9.93 | 3.66 | 1.28 | 6.73 |
| GP2 | F(6)A2 | H3N4F1 | 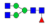 | √ | √ | √ | 1485.5337 | 1488.5556 | 5.98 | 3.02 | 3.92 | 6.28 |
| GP4 | F(6)A2[6]G1 | H4N4F1 | 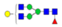 | √ | √ | √ | 1647.5865 | 1650.6084 | 4.16 | 1.81 | 5.87 | 5.56 |
| GP5 | F(6)A2[3]G1 | | | | | | | | | | | |
| × | A2G2 | H5N4 | 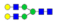 | √ | √ | × | 1663.5814 | 1666.6031 | 8.35 | 2.11 | 11.89 | 9.37 |
| GP3 | F(6)A2B | H3N5F1 | 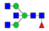 | √ | √ | √ | 1688.613 | 1691.6349 | 3.06 | 1.26 | 1.12 | 5.03 |
| GP9 | F(6)A1G1S1 | H4N3F1Ge1 | 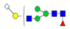 | √ | √ | √ | 1779.6287 | 1782.6507 | 11.07 | 9.94 | 11.16 | 12.37 |
| GP8 | F(6)A2G2 | H5N4F1 | 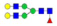 | √ | √ | √ | 1809.6393 | 1812.6612 | 4.54 | 2.12 | 7.85 | 7.04 |
| GP6 | F(6)A2[6]BG1 | H4N5F1 | 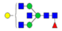 | × | √ | √ | 1850.6659 | 1853.6878 | 0.60 | 4.82 | 2.79 | 4.05 |
| GP10 | F(6)A2[6]G1S1 | H4N4F1Gl1 | 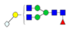 | × | √ | √ | 1936.6662 | 1939.6882 | 1.30 | 0.49 | 9.16 | 6.21 |
| GP11 | F(6)A2[3]G1S1 | | | | | | | | | | | |
| GP10 | F(6)A2[6]G1S1 | H4N4F1Ge1 | | √ | √ | | 1982.7081 | 1985.73 | 5.22 | 2.39 | 6.33 | 6.63 |
| GP11 | F(6)A2[3]G1S1 | H4N4F1Ge1 | 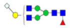 | | | | | | | | | |
| GP12 | A2G2S1 | H5N4Ge1 | 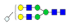 | √ | √ | √ | 1998.703 | 2001.7249 | 9.51 | 5.38 | 5.23 | 7.81 |
| GP13 | F(6)A2G2S1 | H5N4F1Gl1 | 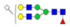 | × | √ | √ | 2098.7191 | 2101.741 | 7.55 | 11.58 | 4.96 | 6.95 |
| GP13 | | H5N4F1Ge1 | 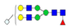 | √ | √ | | 2144.7609 | 2147.7829 | 4.03 | 2.26 | 5.20 | 5.97 |
| GP14 | F(6)A2G2G(3)1S1 | H6N4F1Ge1 | 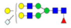 | √ | √ | √ | 2306.8138 | 2309.8357 | 5.96 | 4.77 | 5.04 | 5.49 |
| GP15 | A2G2S2 | H5N4Gl1Ge1 | 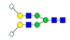 | × | √ | √ | 2287.7828 | 2290.8047 | 3.99 | 9.28 | 0.16 | 9.27 |
| GP15 | | H5N4Ge2 | 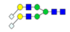 | √ | √ | | 2333.8247 | 2336.8466 | 8.02 | 11.79 | 4.61 | 9.14 |
| GP16 | F(6)A2G2S2 | H5N4F1Gl1Ge1 | 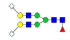 | × | √ | √ | 2433.8407 | 2436.8626 | 17.74 | 8.22 | 1.38 | 9.69 |
| GP16 | | H5N4F1Ge2 | 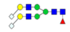 | √ | √ | | 2479.8826 | 2482.9045 | 8.41 | 4.10 | 7.20 | 8.90 |
