## Supplementary material for "Absolute Quantification of Aging-Associated Glycans in IgG for Biological Age Prediction: Insights from Glycomics and Transcriptomics": sTable2-1

**Supplementary Table 2-1:** The genes included in protein N-linked glycosylation (GO:0006487) retrieved from the Mouse Genome Database (MGD).

| MGI Gene/Marker ID | Symbol |
|---|---|
| MGI:1914731 | Alg2 |
| MGI:1913498 | Alg5 |
| MGI:2444031 | Alg6 |
| MGI:2141959 | Alg8 |
| MGI:1924753 | Alg9 |
| MGI:2146159 | Alg10b |
| MGI:2142632 | Alg11 |
| MGI:2385025 | Alg12 |
| MGI:1914824 | Alg13 |
| MGI:1914039 | Alg14 |
| MGI:95705 | B4galt1 |
| MGI:2384987 | B4galt7 |
| MGI:101912 | Dad1 |
| MGI:1194508 | Ddost |
| MGI:1917627 | Derl3 |
| MGI:1914672 | Dhdds |
| MGI:1914093 | Dolpp1 |
| MGI:1196396 | Dpagt1 |
| MGI:1330239 | Dpm1 |
| MGI:1321385 | Entpd5 |
| MGI:108050 | Ext2 |
| MGI:95594 | Fut4 |
| MGI:1858901 | Fut8 |
| MGI:1330859 | Fut9 |
| MGI:95698 | Gfpt1 |
| MGI:1338883 | Gfpt2 |
| MGI:1921748 | Gorasp1 |
| MGI:1913309 | Krtcap2 |
| MGI:1914325 | Magt1 |
| MGI:96973 | Mgat1 |
| MGI:2384966 | Mgat2 |
| MGI:104532 | Mgat3 |
| MGI:2662992 | Mgat4a |
| MGI:2143974 | Mgat4b |
| MGI:1914819 | Mgat4c |
| MGI:1914805 | Mgat4d |
| MGI:1918251 | Mgat4e |
| MGI:3045320 | Mgat4f |
| MGI:894701 | Mgat5 |
| MGI:3606200 | Mgat5b |
| MGI:1924015 | Mlec |

Continued from previous page

| MGI Gene/Marker ID | Symbol |
|---|---|
| MGI:1929872 | Mogs |
| MGI:1196365 | Nus1 |
| MGI:1914945 | Ost4 |
| MGI:1913607 | Ostc |
| MGI:3839961 | Pate6 |
| MGI:97566 | Pgm3 |
| MGI:1353418 | Pmm1 |
| MGI:1859214 | Pmm2 |
| MGI:3607791 | Rft1 |
| MGI:98084 | Rpn1 |
| MGI:98085 | Rpn2 |
| MGI:891971 | Serpina1a |
| MGI:891970 | Serpina1b |
| MGI:1914797 | Slc39a8 |
| MGI:1930252 | Srd5a3 |
| MGI:108470 | St6gal1 |
| MGI:105124 | Stt3a |
| MGI:1915542 | Stt3b |
| MGI:894407 | Tmem165 |
| MGI:1916288 | Tmem258 |
| MGI:1933134 | Tusc3 |
| MGI:1926245 | Ube2j1 |
| MGI:2443162 | Uggt1 |
| MGI:1913685 | Uggt2 |
