## Supplementary material for "Absolute Quantification of Aging-Associated Glycans in IgG for Biological Age Prediction: Insights from Glycomics and Transcriptomics": sTable2-2

**Supplementary Table 2-2**: The candidate genes identified by previous GWAS

| HGNC. symbol | IgG N-glycan Trait | Functional.validation.in..vitro | Reference.study | MGI. symbol |
|---|---|---|---|---|
| RUNX3 | G0 | Negative Correlation with G | ( Lauc,G et al., 2023; Mijakovac et.al.,2021; Shadrina, A.S., 2021) | Runx3 |
| KIF3C | G0 | ns | ( Lauc,G et al., 2023) | Kif3c |
| NFKB1 | G2 | ns | ( Lauc,G et al., 2023) | Nfkb1 |
| MANBA | G2 | Negative Correlation with G | ( Lauc,G et al., 2023) | Manba |
| HLA region | G0 | | ( Lauc,G et al., 2023) | |
| EEF1A1 | G2 | Negative Correlation with G | ( Lauc,G et al., 2023) | Eef1a1 |
| MTO1 | G2 | | ( Lauc,G et al., 2023) | Mto1 |
| HIVEP2 | G2;F | Positive Correlation with G but no statistically significant change in HIVEP2 transcript level | ( Lauc,G et al., 2023; Shadrina, A.S., 2021) | Hivep2 |
| ABCF2 | G0 | | ( Lauc,G et al., 2023; Shadrina, A.S., 2021) | Abcf2 |
| CHPF2 | G0 | | ( Lauc,G et al., 2023) | Chpf2 |
| SMARCD3 | G0 | | ( Lauc,G et al., 2023) | Smarcd3 |
| UBR5 | G0 | | ( Lauc,G et al., 2023) | Ubr5 |
| RRM2B | G0 | | ( Lauc,G et al., 2023) | Rrm2b |
| ODF1 | G0;B | | ( Lauc,G et al., 2023; Shadrina, A.S., 2021) | Odf1 |
| KB-1980E6.3 | G0 | | ( Lauc,G et al., 2023) | |
| B4GALT1 | G2 | | ( Lauc,G et al., 2023) | B4galt1 |
| OVOL1 | G1 | | ( Lauc,G et al., 2023; Shadrina, A.S., 2021) | Ovol1 |
| AP5B1 | G1 | | ( Lauc,G et al., 2023) | Ap5b1 |
| TNFRSF13B | G1 | Negative Correlation with G0 | ( Lauc,G et al., 2023; Shadrina, A.S., 2021) | Tnfrsf13b |
| ARHGAP27 | G1 | | ( Lauc,G et al., 2023) | Arhgap27 |

Continued from previous page

| HGNC. symbol | IgG N-glycan Trait | Functional.validation.in..vitro | Reference.study | MGI. symbol |
|---|---|---|---|---|
| CRHR1 | G1 | | ( Lauc,G et al., 2023) | |
| SPPL2C | G1 | | ( Lauc,G et al., 2023) | Sppl2c |
| MAPT | G1 | | ( Lauc,G et al., 2023; Shadrina, A.S., 2021) | Mapt |
| KANLS1 | G0 | | ( Lauc,G et al., 2023) | |
| ARL17B | G0 | | ( Lauc,G et al., 2023) | Arf2 |
| LRRC37A | G0 | | ( Lauc,G et al., 2023) | Lrrc37a |
| LRRC37A | G0 | | ( Lauc,G et al., 2023) | Gm884 |
| NSF | G0 | | ( Lauc,G et al., 2023) | Nsf |
| WNT3 | G0 | | ( Lauc,G et al., 2023) | Wnt3 |
| TBKBP1 | G2 | | ( Lauc,G et al., 2023) | Tbkbp1 |
| TBX21 | G2 | | ( Lauc,G et al., 2023; Shadrina, A.S., 2021) | Tbx21 |
| BZRAP1 | G2 | | ( Lauc,G et al., 2023) | |
| SUPT4H1 | G2 | | ( Lauc,G et al., 2023) | Supt4a |
| RAD5C1 | G2 | | ( Lauc,G et al., 2023) | |
| AZI1 | G2 | | ( Lauc,G et al., 2023) | |
| ENTHD2 | G2 | | ( Lauc,G et al., 2023) | |
| SLC38A10 | G2 | ns | ( Lauc,G et al., 2023) | Slc38a10 |
| C17orf89 | G2 | | ( Lauc,G et al., 2023) | |
| RUNX1 | | ns | (Mijakovac et.al.,2021; Shadrina, A.S., 2021) | Runx1 |
| SPINK4 | G | Negative Correlation with G | (Mijakovac et.al.,2021) | Spink4 |
| SPPL3 | G | Positive Correlation with G | (Mijakovac et.al.,2022) | Sppl3 |
| IKZF1 | F; G | | ( Lauc,G et al., 2013; Shadrina, A.S., 2021) | Ikzf1 |
| IL6ST | G | | ( Lauc,G et al., 2013) | |
| ANKRD55 | G | | ( Lauc,G et al., 2013) | Ankrd55 |
| SMARCB1 | B | | ( Lauc,G et al., 2013;Shadrina, A.S., 2021 ) | Smarcb1 |
| DERL3 | B | | ( Lauc,G et al., 2013) | Derl3 |
| SMYD3 | G | | (Shadrina, A.S., 2021) | Smyd3 |
| ASXL2 | B | | (Shadrina, A.S., 2021) | Asxl2 |
| CHST2 | | | (Shadrina, A.S., 2021) | Chst2 |

Continued from previous page

| HGNC. symbol | IgG N-glycan Trait | Functional.validation.in..vitro | Reference.study | MGI. symbol |
|---|---|---|---|---|
| SLC9A9 | | | (Shadrina, A.S., 2021) | Slc9a9 |
| RNF168 | | | (Shadrina, A.S., 2021) | Rnf168 |
| KDELR2 | F | | (Shadrina, A.S., 2021) | Kdelr2 |
| COG7 | F | | (Shadrina, A.S., 2021) | Cog7 |
| MEF2B | F | | (Shadrina, A.S., 2021) | Mef2b |
| BORCS8-MEF2B | F | | (Shadrina, A.S., 2021) | |
| MGME1 | S | | (Shadrina, A.S., 2021) | Mgme1 |
| BANF2 | S | | (Shadrina, A.S., 2021) | Banf2 |
| SNXS | S | | (Shadrina, A.S., 2021) | |
| MIR4425 | G | | (Shadrina, A.S., 2021) | |
| ST6GAL1 | G | | (Shadrina, A.S., 2021) | St6gal1 |
| ELL2 | | | (Shadrina, A.S., 2021) | Ell2 |
| IRF1-AS1 | | | (Shadrina, A.S., 2021) | |
| IRF1 | | | (Shadrina, A.S., 2021) | Irf1 |
| SLC22A5 | | | (Shadrina, A.S., 2021) | Slc22a5 |
| SLC22A5 | | | (Shadrina, A.S., 2021) | Slc22a21 |
| MICB-DT | F | | (Shadrina, A.S., 2021) | |
| MICB | F | | (Shadrina, A.S., 2021) | Mill1 |
| MICB | F | | (Shadrina, A.S., 2021) | Mill2 |
| HCG26 | F | | (Shadrina, A.S., 2021) | |
| TXINB | F | | (Shadrina, A.S., 2021) | |
| BAGALT1 | F | | (Shadrina, A.S., 2021) | |
| NXPE4 | | | (Shadrina, A.S., 2021) | Nxpe4 |
| NXPE2 | | | (Shadrina, A.S., 2021) | Nxpe2 |
| MIR4708 | | | (Shadrina, A.S., 2021) | |
| FUT8 | | | (Shadrina, A.S., 2021) | Fut8 |
| TMEM121 | | | (Shadrina, A.S., 2021) | Tmem121 |
| IKZF3 | G1 | | (Shadrina, A.S., 2021) | Ikzf3 |
| GSDMB | G1 | | (Shadrina, A.S., 2021) | |
| ORMDL3 | G1 | | (Shadrina, A.S., 2021) | Ormdl3 |
| IRRC3C | G1 | | (Shadrina, A.S., 2021) | |
| ZPBP2 | G1 | | (Shadrina, A.S., 2021) | Zpbp2 |
| CEP131 | F | | (Shadrina, A.S., 2021) | Cep131 |
| FUT6 | | | (Shadrina, A.S., 2021) | |
| RUNX1-IT1 | | | (Shadrina, A.S., 2021) | |
| TAB1 | | | (Shadrina, A.S., 2021) | Tab1 |
| MGAT3 | | | (Shadrina, A.S., 2021) | Mgat3 |

Note：

Continued from previous page

(Lauc,G et al., 2023) Frkatović-Hodžić, A., Mijakovac, A., Miškec, K., Nostaeva, A., Sharapov, S.Z., Landini, A., Haller, T., Akker, E.V.D., Sharma, S., Cuadrat, R.R.C., Mangino, M., Li, Y., Keser, T., Rudman, N., Štambuk, T., Pučić-Baković, M., Trbojević-Akmačić, I., Gudelj, I., Štambuk, J., Pribić, T., Radovani, B., Tominac, P., Fischer, K., Beekman, M., Wuhrer, M., Gieger, C., Schulze, M.B., Wittenbecher, C., Polasek, O., Hayward, C., Wilson, J.F., Spector, T.D., Köttgen, A., Vučković, F., Aulchenko, Y.S., Vojta, A., Krištić, J., Klarić, L., Zoldoš, V., Lauc, G., 2023. Mapping of the gene network that regulates glycan clock of ageing. Aging (Albany NY) 15, 14509-14552.

(Lauc,G et al., 2013) Lauc, G., Huffman, J.E., Pučić, M., Zgaga, L., Adamczyk, B., Mužinić, A., Novokmet, M., Polašek, O., Gornik, O., Krištić, J., Keser, T., Vitart, V., Scheijen, B., Uh, H.W., Molokhia, M., Patrick, A.L., McKeigue, P., Kolčić, I., Lukić, I.K., Swann, O., van Leeuwen, F.N., Ruhaak, L.R., Houwing-Duistermaat, J.J., Slagboom, P.E., Beekman, M., de Craen, A.J., Deelder, A.M., Zeng, Q., Wang, W., Hastie, N.D., Gyllensten, U., Wilson, J.F., Wuhrer, M., Wright, A.F., Rudd, P.M., Hayward, C., Aulchenko, Y., Campbell, H., Rudan, I., 2013. Loci associated with N-glycosylation of human immunoglobulin G show pleiotropy with autoimmune diseases and haematological cancers. PLoS Genet 9, e1003225.

(Mijakovac et.al.,2021) Mijakovac, A., Jurić, J., Kohrt, W.M., Krištić, J., Kifer, D., Gavin, K.M., Miškec, K., Frkatović, A., Vučković, F., Pezer, M., Vojta, A., Nigrović, P.A., Zoldoš, V., Lauc, G., Effects of Estradiol on Immunoglobulin G Glycosylation: Mapping of the Downstream Signaling Mechanism.

(Mijakovac et.al.,2022) Mijakovac, A., Miškec, K., Krištić, J., Vičić Bočkor, V., Tadić, V., Bošković, M., Lauc, G., Zoldoš, V., Vojta, A., 2022. A Transient Expression System with Stably Integrated CRISPR-dCas9 Fusions for Regulation of Genes Involved in Immunoglobulin G Glycosylation. Crispr j 5, 237-253.

(Shadrina, A.S., 2021) Shadrina, A.S., Zlobin, A.S., Zaytseva, O.O., Klarić, L., Sharapov, S.Z., D Pakhomov, E., Perola, M., Esko, T., Hayward, C., Wilson, J.F., Lauc, G., Aulchenko, Y.S., Tsepilov, Y.A., 2021. Multivariate genome-wide analysis of immunoglobulin G N-glycosylation identifies new loci pleiotropic with immune function. Human Molecular Genetics 30, 1259-1270.

ns No Significance
